# Transfer rate of enveloped and non-enveloped viruses between fingerpads and surfaces

**DOI:** 10.1101/2021.06.22.449538

**Authors:** Claire E. Anderson, Alexandria B. Boehm

## Abstract

Fomites can represent a reservoir for pathogens, which may be subsequently transferred from surfaces to skin. In this study we aim to understand how different factors (including virus type, surface type, time since last handwash, and direction of transfer) affect virus transfer rates, defined as the fraction of virus transferred, between fingerpads and fomites. To determine this, 360 transfer events were performed with 20 volunteers using Phi6 (a surrogate for enveloped viruses) and MS2 (a surrogate for non-enveloped viruses), and three clean surfaces (stainless steel, painted wood, and plastic). Considering all transfer events (all surfaces and both transfer directions combined), the mean transfer rates of Phi6 and MS2 were 0.17 and 0.26, respectively. Transfer of MS2 was significantly higher than Phi6 (P<0.05). Surface type was a significant factor that affected the transfer rate of Phi6: Phi6 is more easily transferred to and from stainless steel and plastic than to and from painted wood. Direction of transfer was a significant factor affecting MS2 transfer rates: MS2 is more easily transferred from surfaces to fingerpads than from fingerpads to surfaces. Data from these virus transfer events, and subsequent transfer rate distributions, provide information which can be used to refine quantitative microbial risk assessments. This study is the first to provide a large-scale data set of transfer events with a surrogate for enveloped viruses, which extends the reach of the study to the role of fomites in the transmission of human enveloped viruses like influenza and SARS-CoV-2.

**Importance:** This study created the first large-scale data set for the transfer of enveloped viruses between skin and surfaces. The data set produced by this study provides information on modelling the distribution of enveloped and non-enveloped virus transfer rates, which can aid in the implementation of risk assessment models in the future. Additionally, enveloped and non enveloped viruses were applied to experimental surfaces in an equivalent matrix to avoid matrix effects, so results between different viral species can be directly compared without confounding effects of different matrices. Our results indicating how virus type, surface type, time since last handwash, and direction of transfer affect virus transfer rates can be used in decision-making processes to lower the risk of viral infection from transmission through fomites.

## Introduction

Viruses are deposited in the environment when fluids (mucus, saliva, urine, feces) containing high viral titer are released from an infected individual (1, 2). Humans can come into contact with viruses when they consume or recreate in virus-contaminated water, eat contaminated food, breathe contaminated air, or touch contaminated fomites. When transmission of a virus occurs via an environmental intermediary, the transmission is referred to as “indirect”. It is well-understood that indirect transmission is important for many viruses including those that cause diarrheal illness, influenza, COVID-19, and measles (2–7). While it is well known that fomite-mediated transmission is an important pathway for many diseases, several studies have emphasized the need for more information about inactivation rates, transfer rates, and pathogen shedding in order to develop accurate exposure and risk models (2, 3, 6, 8, 9).

Transmission of viruses via contaminated fomites requires multiple steps (Figure 1). First, a susceptible individual must come into the contact with the surface. Second, viruses are transferred between the fomite and the susceptible individual. Third, the virus transferred via touch is transmitted to the individual. The last step may require an additional transfer event from the part of the body that touched the fomite to another part of the body where infection occurs (sometimes referred to as self-inoculation). Whether the transmission event results in infection depends on the biology of the virus and the immune system of the individual. Infected individuals can also deposit viruses onto fomites via touch if there is virus present on their body, thereby contaminating fomites with viruses. In the present study, we are particularly focused on the transfer of viruses to and from skin and fomites.

**Figure 1.**
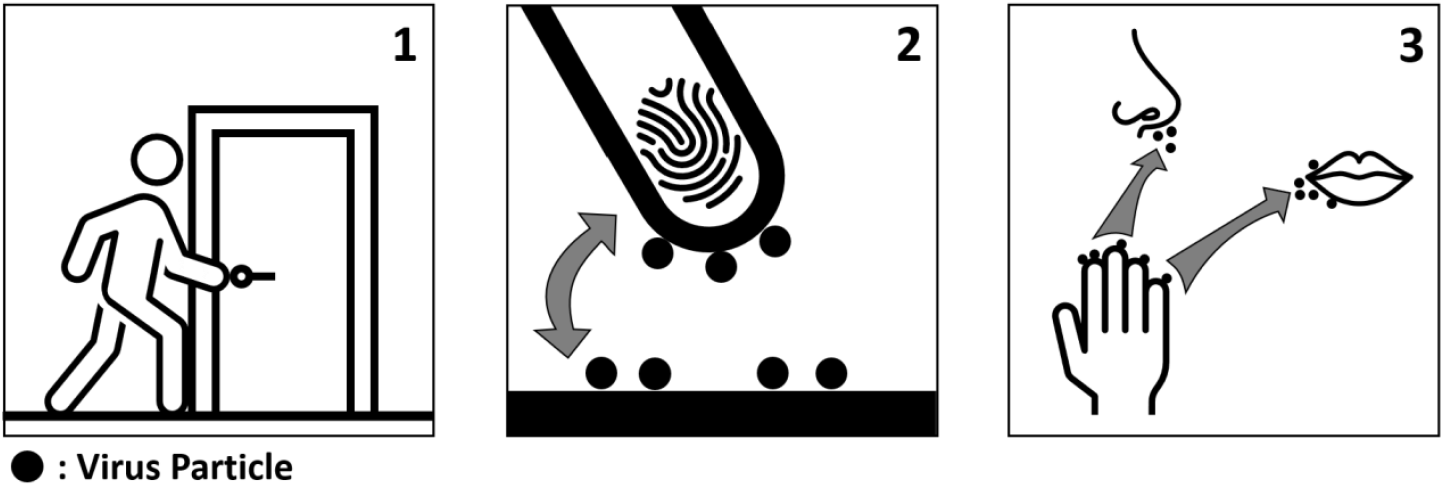
Pathway of transmission from contaminated fomites. First the individual contacts a surface, on which the individual picks up or deposits invective virus particles. Lastly, the individual transfers the infective virus from their hand to an area of their body where infection occurs, or to an additional surface.

Six studies have characterized the transfer of viruses to and from skin and inanimate surfaces and they have been primarily been undertaken using non-enveloped viruses (Table 1) (1, 5, 10–13). This collection of studies includes virus transfer studies that explicitly quantified transfer rates between human skin and non-food surfaces (14) (see Table 1 of Zhao et al. (14) for a complete list of all virus transfer studies). These studies quantified transfer of MS2 (1, 10), poliovirus 1 (1), human parainfluenza virus (5), rhinovirus (5, 13), ϕX174 (10), fr (10), rotavirus (11), and hepatitis A (12). Of these viruses, only human parainfluenza is enveloped. When investigating human parainfluenza virus transfer between skin and surfaces, Ansari et al. (5) found that the virus inactivated quickly and therefore it was impossible to quantify the transfer rate. Although experimental variables such as humidity, surface type, and virus type vary for each study, aspects of the experimental procedures remain relatively consistent for each study. Four of the six studies (1, 5, 11, 12) have an inoculation volume of 10 μL, a contact time of 5-10 seconds, 20-30 minutes of inoculum dry time, and a contact pressure of 1 kg/cm^2^. All 6 studies quantify virus transfer rate which is defined as the fraction of virus transferred upon contact. Julian et al. (10) defines transfer rate as the PFU recovered from the non-inoculated surface over the total PFU recovered from both surfaces. One of the studies investigated transfer rates between both porous and nonporous fomites (1), while the rest studied only nonporous fomites. Across all 6 studies, transfer rate varied between <0.02 to 0.80 for nonporous surfaces (1, 5, 10–13). The study investigating porous surfaces found a transfer rate of <0.07 (1).

**Table 1.** A summary of previous studies investigating viral transfer rates. Included in the table are various experimental variables, including virus species, inanimate surface, inoculation volume, contact time, inoculum dry time, and contact pressure.

| Reference | Virus Species | Surfaces | Inoculation Volume | Contact Time | Inoculum Dry Time | Contact Pressure | Transfer Rate |
| --- | --- | --- | --- | --- | --- | --- | --- |
| Reed, 1975 (13) | Rhinovirus | Plastic pen<br>Table<br>Stainless steel | 2-5.5 µL | 15 s | 2-10 min | Not recorded | Mean up to 0.46 |
| Ansari et al., 1988 (11) | Rotavirus | Stainless steel | 10 µL | 10 s | 20 min | 1 kg/cm<sup>2</sup> | Mean up to 16.8 |
| Ansari et al., 1991 (5) | Human parainfluenza virus 3<br>Rhinovirus 14 | Stainless Steel | 10 µL | 5 s | 20 min | 1.0 kg/cm<sup>2</sup> | Mean up to 0.02 |
| Mbithi et al., 1992 (12) | Hepatitis A | Stainless Steel | 10 µL | 10 s | 20 min | 0.2 kg/cm<sup>2</sup> to<br>1 kg/cm<sup>2</sup> | Mean up to 0.27 |
| Julian et al., 2010 (10) | MS2<br>φX174<br>fr | Glass | 5 µL | 10 s | 10 min | 0.25 kg/cm<sup>2</sup> | Mean of 0.23 |
| Lopez et al., 2013 (1) | MS2<br>Poliovirus 1 | Acrylic<br>Glass<br>Ceramic tile<br>Laminate<br>Stainless steel<br>Granite<br>Cotton<br>Polyester<br>Paper Currency | 10 µL | 10 s | 30 min | 1.0 kg/cm<sup>2</sup> | Nonporous Surfaces:<br>Mean up to 0.80<br>Porous Surfaces: Mean<br>of <0.07 |
| This Study | MS2<br>Phi6 | Plastic<br>Stainless steel<br>Painted Wood | 10 µL | 10 s | Up to 30<br>min | 0.25 kg/cm<sup>2</sup> | Mean up to 0.26 |

Zhao et al. (14) provide a mechanistic model of transfer rates between surfaces. Their model considers the physical and chemical mechanisms that control transfer. Their model suggests touch force, microbial diameter, inoculation volume, touch number by the same individual, rubbing, and humidity have a positive correlation with virus transfer. They also suggest donor roughness, touch number by different individuals, surface hardness, temperature, surface inoculation area, and surface touching area are negatively correlated to virus transfer.

There is presently no experimental data on transfer of enveloped viruses between skin and surfaces, so this study sought to fill that knowledge gap. We documented the transfer rate of enveloped and non-enveloped viruses between various surfaces and fingertips using human volunteers and 360 transfer events, creating the first large-scale data set for enveloped viruses. The data set produced by this study provides information on modelling the distribution of enveloped and non-enveloped virus transfer rates, which can aids in the implementation of risk assessment models in the future (8, 15–17).

We also investigated how virus type, surface type, time since last handwash, and direction of transfer affect virus transfer rates. The choice of variables is informed by results of previous studies and the model developed by Zhao et al. (14). Enveloped and non-enveloped viruses were applied to experimental surfaces in an equivalent matrix in order to avoid matrix effects, so the results obtained with different viral species can be directly compared without confounding effects of different matrices.

The enveloped virus used in this study is Phi6. Phi6 has a dsRNA genome and is spherical in shape with ~80-100 nm diameter; the protein nucleocapsid is surrounded by a lipid membrane and thus, it serves as a non-pathogenic, biosafety-level 1 bacteriophage surrogate for enveloped human pathogenic viruses, such as influenza, SARS-CoV-2, and Ebola. The non-enveloped virus used in this study is MS2. MS2 has an ssRNA genome and has an icosahedral protein shell ~27 nm in diameter. MS2 similarly acts as a biosafety level-1 bacteriophage surrogate for non-enveloped human pathogenic viruses such as norovirus and enteroviruses. Phi6 and MS2 have been previously applied to hands to model pathogenic viruses (10, 18–20).

## Results

### Experimental Conditions

A total of 20 volunteers participated in the study. They ranged in age from 22 to 58 years, with the median age being 26. Volunteer hand length ranged from 16.2 cm to 21.9 cm, with the median length being 19.3 cm. Volunteer hand breadth ranged from 7.3 cm to 10 cm, with a median breadth of 8.1 cm. Temperature of the laboratory throughout the study ranged from 20.8°C to 21.9°C, with a median temperature of 21.7°C. Relative humidity during the study ranged from 13% to 74%, with a median value of 58%. Full temperature and humidity data are available in the SI.

### Transfer Rate Distributions

All negative controls had 0 PFU and all viral stock concentrations had an expected number of PFU/mL.

The fraction of virus transferred (*f*) was determined for 360 transfer events for the two viruses. Out of the 360 transfer events for Phi6, all three dilutions plated were TNTC 8 times. All three dilutions were lower than the detection limit 38 times. As a result, 46 transfer events were removed from the data set for Phi6, leaving 314. Out of the 360 transfer events for MS2, there were no instances where all dilutions exceeded the limit of detection. The three dilutions were lower than the detection limit 4 times for MS2. As a result, 4 transfer events were removed from the data set for MS2, leaving 356. The instances where the transfer rate was irrecoverable for Phi6 and MS2 are not limited to a single surface, time since last handwash, or direction of transfer. The instances also make up less than 7% of the total data, and therefore are not anticipated to affect the overall distribution of the data. More information about these instances of irrecoverable transfer rates can be found in Table 2.

**Table 2.** Descriptive statistics for the transfer rate of Phi6 and MS2. Statistics are broken down by virus type, surface, time since last handwash, and direction of transfer. Included in the statistics are the number of trials for each condition, the mean, the median, and the standard deviation of the transfer rate.

| Phi6 |  |  |  |  |  |  |
| --- | --- | --- | --- | --- | --- | --- |
| Surface | Time since last handwash | Direction of transfer | # of Trials | Mean | Median | Standard Deviation |
| Stainless steel | 1 hour | Surface to fingerpad | 30 | 0.23 | 0.19 | 0.19 |
|  |  | Fingerpad to surface | 25 | 0.18 | 0.16 | 0.20 |
|  | 0 hour | Surface to fingerpad | 30 | 0.20 | 0.17 | 0.15 |
|  |  | Fingerpad to surface | 22 | 0.22 | 0.21 | 0.15 |
| Plastic | 1 hour | Surface to fingerpad | 30 | 0.28 | 0.22 | 0.23 |
|  |  | Fingerpad to surface | 22 | 0.17 | 0.09 | 0.19 |
|  | 0 hour | Surface to fingerpad | 30 | 0.22 | 0.21 | 0.14 |
|  |  | Fingerpad to surface | 22 | 0.15 | 0.11 | 0.12 |
| Wood | 1 hour | Surface to fingerpad | 28 | 0.05 | 0.01 | 0.07 |
|  |  | Fingerpad to surface | 26 | 0.13 | 0.09 | 0.14 |
|  | 0 hour | Surface to fingerpad | 27 | 0.08 | 0.03 | 0.10 |
|  |  | Fingerpad to surface | 22 | 0.07 | 0.05 | 0.06 |
| MS2 |  |  |  |  |  |  |
| Surface | Time since last handwash | Direction of transfer | # of Trials | Mean | Median | Standard Deviation |
| Stainless steel | 1 hour | Surface to fingerpad | 30 | 0.34 | 0.33 | 0.12 |
|  |  | Fingerpad to surface | 30 | 0.13 | 0.08 | 0.12 |
|  | 0 hour | Surface to fingerpad | 30 | 0.37 | 0.37 | 0.12 |
|  |  | Fingerpad to surface | 30 | 0.18 | 0.13 | 0.17 |
| Plastic | 1 hour | Surface to fingerpad | 30 | 0.37 | 0.33 | 0.14 |
|  |  | Fingerpad to surface | 30 | 0.16 | 0.11 | 0.16 |
|  | 0 hour | Surface to fingerpad | 30 | 0.40 | 0.37 | 0.18 |
|  |  | Fingerpad to surface | 29 | 0.15 | 0.11 | 0.17 |
| Wood | 1 hour | Surface to fingerpad | 30 | 0.30 | 0.29 | 0.18 |
|  |  | Fingerpad to surface | 28 | 0.22 | 0.17 | 0.17 |
|  | 0 hour | Surface to fingerpad | 30 | 0.33 | 0.29 | 0.20 |
|  |  | Fingerpad to surface | 29 | 0.21 | 0.19 | 0.18 |

The mean transfer rate for Phi6 was 0.17, while the median was 0.12 and the standard deviation was 0.17. For MS2, the mean transfer rate was 0.26, the median was 0.25, and the standard deviation was 0.18 (Figures 3 and 4). The respective means, medians, and standard deviations of the transfer rate based on the variables investigated (virus type, surface type, time since last handwash, and direction of transfer) can be found in Table 2 and Figure 3.

**Figure 2.**
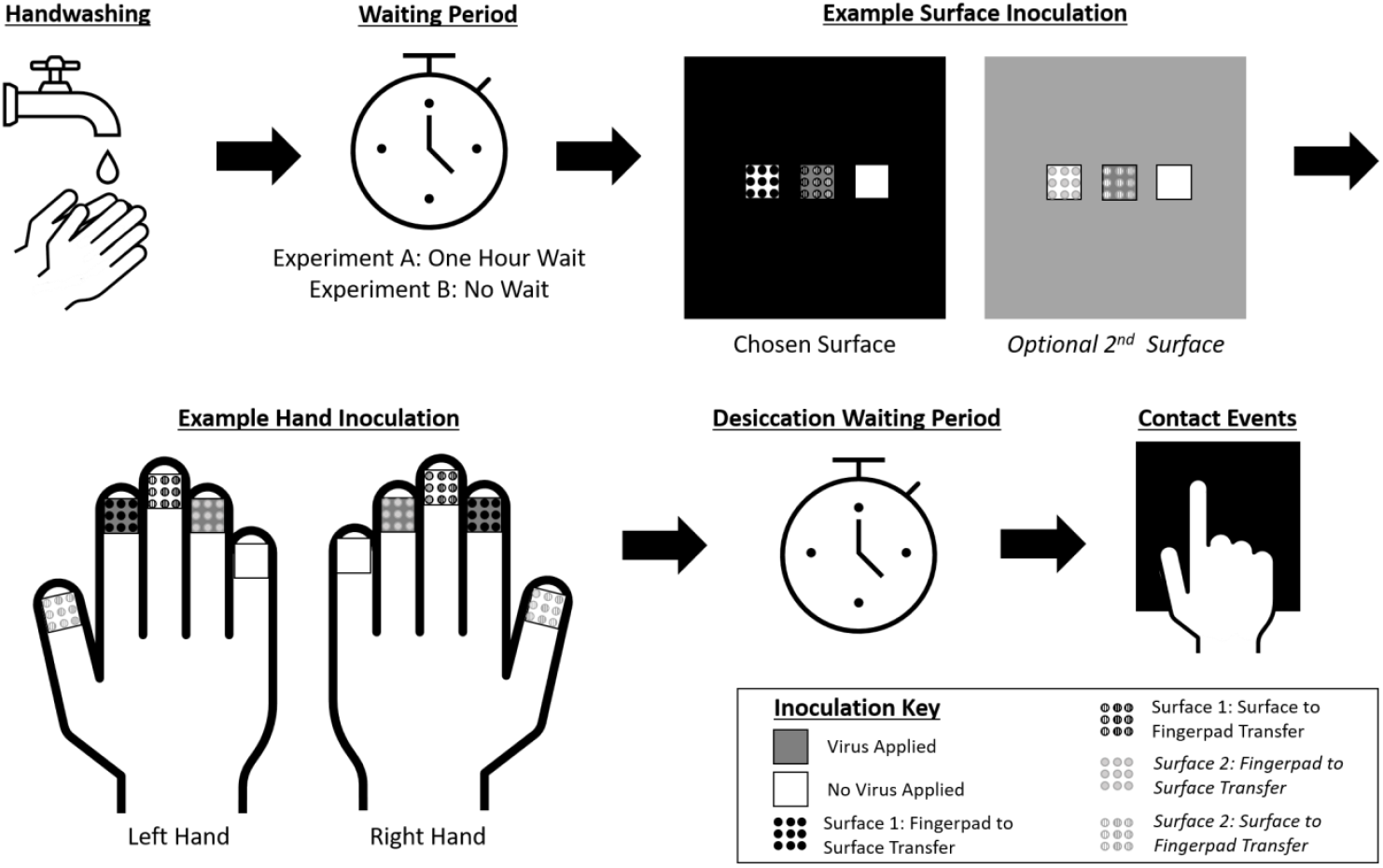
Outline of experimental procedure up until the contact events. The procedure outlines the initial hand washing step, followed by the wait time for Experiment A or B, an example of surface inoculation, an example of hand inoculation, the wait time for the inoculum to dry, and the contact events. In the example surfaces inoculation, the leftmost square represents where the virus was not applied, but where the transfer from the fingerpad to the surface would occur. The middle square represents where the virus was applied (indicating transfer from the surface to the fingerpad) and the rightmost is the control. In the example hand inoculation, the hands are duplicates. On the middle finger and thumb there is no virus applied, but represents where the transfer from the chosen surface and 2^nd^ surface to the fingerpad will occur, respectively. On the index finger and ring finger is where there is virus applied, and where transfer between the finger and the respective surfaces will occur. The pinky is a control.

**Figure 3.**
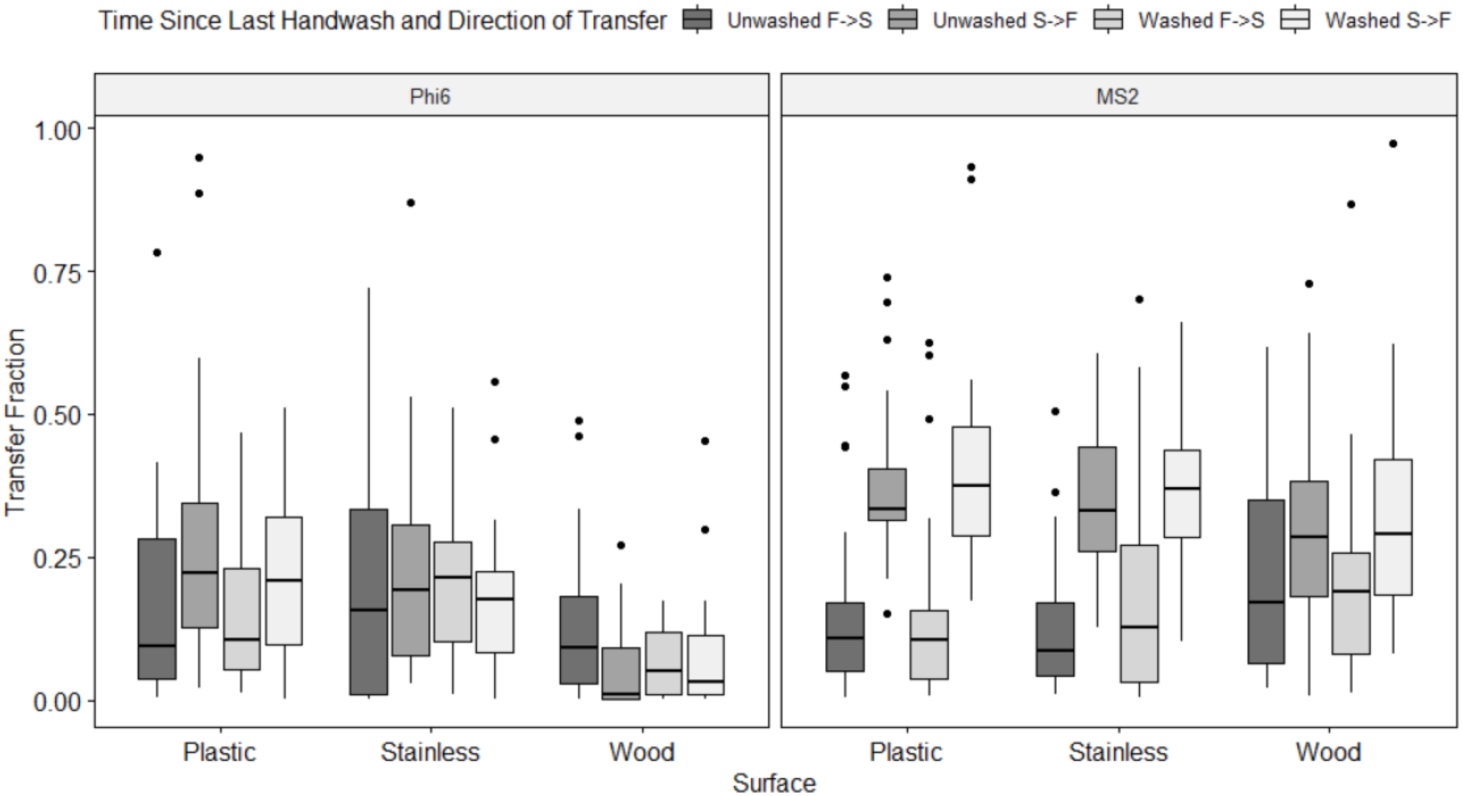
Boxplots of transfer rates for different surfaces. The upper and lower whiskers show the maximum and minimum values, respectively (excluding outliers defined by the interquartile range criterion). The lower and upper edges of the represent the lower and upper quartile, respectively. The horizontal line within the box indicates the median. The points beyond the whiskers represent outliers. The data are broken down by ‘virus type’, ‘surface type’, ‘time since last handwash’, and ‘direction of transfer’. “Unwashed” represents 1 hour since last handwash and “washed” represents 0 hour since last handwash. “F->S” represents fingerpad to surface transfer and “S->F” represents surface to fingerpad transfer.

**Figure 4.**
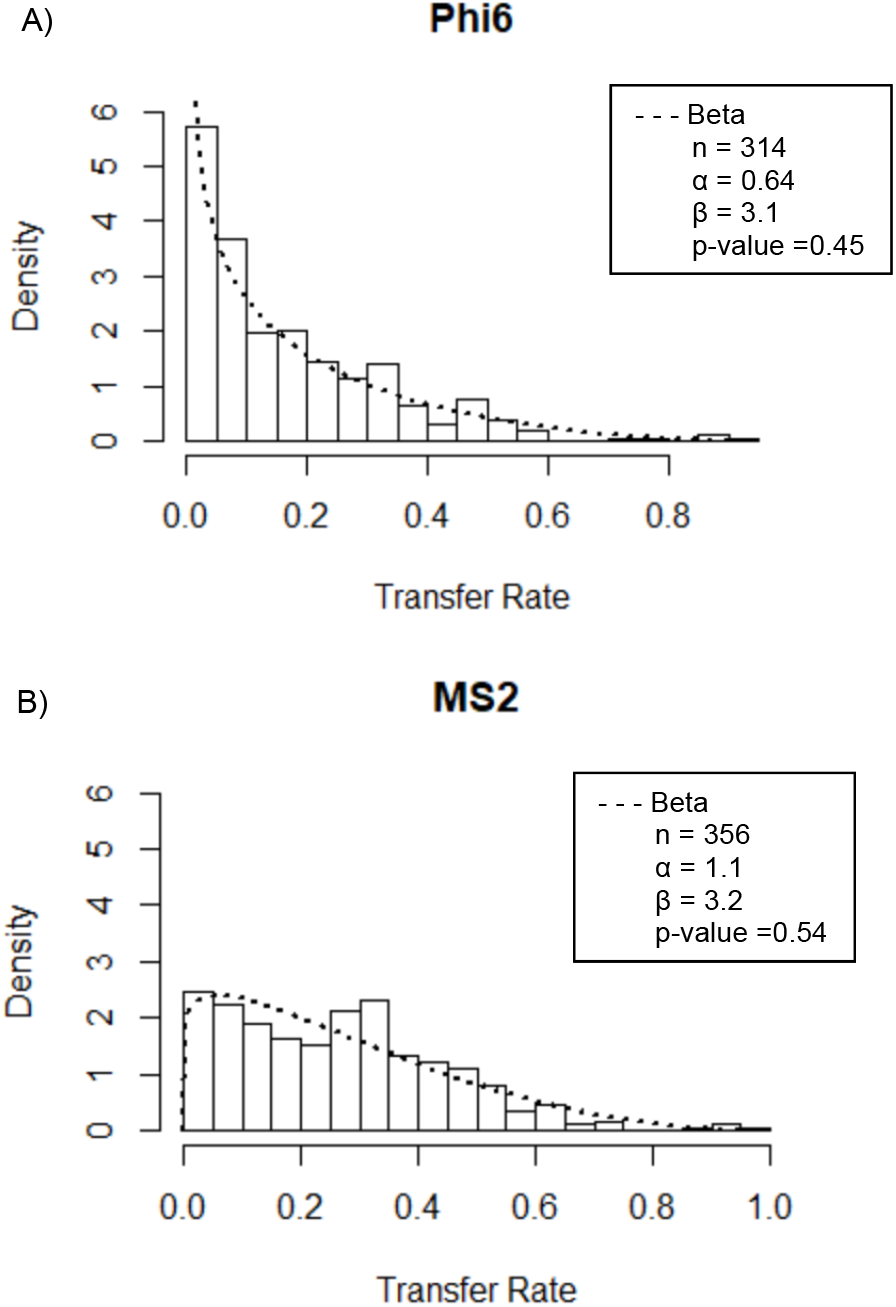
Phi6 (A) and MS2 (B) histogram distributions, overlayed with probability distribution functions. The functions used to model the data are beta (- - -). The alpha and beta shape parameters, as well as the goodness of fit p-value, are also shown.

Several distributions (including normal, lognormal, exponential, geometric, and beta) were fit to the data and the goodness of fit for each was tested through a Kolomogov-Smirnoff test, comparing the log-likelihood, and comparing the Akaike’s Information Criterion (AIC). Overlayed on the histogram in Figure 4 is the distribution that best fit the data of the distributions tested, along with the distribution parameters. In the case of both virus type, beta distributions fit the data best. For each virus, the beta distribution had the highest log-likelihood estimate, the lowest AIC, and a p-value greater than 0.05. Although the normal distribution fit the data well (a p-value of 0.46 and 0.54 for Phi6 and MS2, respectively), it was not used because it included the possibility of negative transfer fraction values, which are physically unrealistic.

### Significant Factors Controlling Transfer Rate

An n-way ANOVA on the complete data set indicates that ‘virus type’ (P<0.001), ‘surface type’ (P<0.001), and ‘direction of transfer’ (P<0.001) are significant factors in controlling transfer. An ANOVA is justified for analyzing these data as the Kolmogov-Smirnoff tested suggested the data could be reasonable approximated as normal. The ‘time since last handwash’ factor was not significant in the model (P=0.87). In terms of interactions between variables, significant two-way interactions were found between the ‘virus type’ and ‘surface type’, the ‘virus type’ and ‘time since last handwash’, and ‘surface-type’ and ‘time since last handwash’. The remaining unlisted interactions were not statistically significant. To parse through these interaction terms, two three-way ANOVAs were performed with Phi6 and MS2 as the dependent variables, separately.

A three-way ANOVA performed with Phi6 transfer rate as the dependent variable indicates that surface type is significant (P<0.001). The post-hoc test shows that there are differences between wood and plastic (mean difference between wood and plastic = −0.13, P<0.001) and wood and stainless steel (mean difference between wood and stainless steel = −0.12, P<0.001), but no difference between stainless steel and plastic (P=0.97). Direction of transfer (P=0.16) and time since last handwash (P=0.24) are not significant factors in the model. There is no statistically significant three-way interaction between ‘ surface type’, ‘ direction of transfer’, or ‘time since last handwash’ (P=0.14). In terms of possible two-way interactions, the only significant interaction occurs between ‘surface type’ and ‘direction of transfer’ (P=0.014); the direction of transfer was found to only significantly impact the transfer rate between fingerpads and plastic (mean difference between finger to plastic transfer and plastic to finger transfer = −0.09).

A separate three-way ANOVA performed for all MS2 data indicates that direction of transfer is the only significant variable (P<0.001). The post-hoc test shows that the mean difference between fingerpad to surface transfer and surface to fingerpad transfer is −0.18. Surface type (P=0.71) and time since last handwash (P=0.23) were not found to be significant. Similarly to Phi6, there is no statistically significant three-way interaction between surface type, direction of transfer, or time since last handwash (P=0.73). The only significant two-way interaction occurs between the surface type and direction of transfer (P=0.003). The direction of transfer significantly effects the transfer from all three surfaces, with a higher fraction transferred from surfaces to fingerpads for all surface types (a mean difference of 0.23 for plastic, 0.21 for stainless steel, and 0.10 for wood).

## Discussion

Both enveloped and non-enveloped viruses are readily transferred between fomites and fingertips with transfer rates of 0.22, on average. This implies that a transfer of 22% of viruses on a surface to a fingerpad should be expected. Whether or not this transfer would result in a risk of fomite-mediated infection would depend on number of infectious viruses contacted by the fingertip, the efficiency of self-inoculation (i.e., transfer of virus from fingertip to the mouth, nasal cavity, or other bodily location where infection may occur), the infectious dose of the virus, and the susceptibility of the individual.

The transfer rates reported in this study for MS2 and Phi6 are similar to virus transfers reported by others (1, 11, 12, 21). Specifically, the MS2 mean transfer rate of 0.26 is comparable to the MS2 mean transfer rate of 0.22 between fingertips and glass reported by Julian et al. (10) who used similar methods as those used herein. Previous work reported viral transfer rates between skin and fomites to range between 0.16 and 0.65 for non-porous surfaces (1, 10–12, 21). The higher values in this range were obtained using greater contact pressure and a shorter desiccation time for viral suspensions (1, 10, 12). According to a physical-chemical model of skin-surface microbial transfer (14), greater contact pressure will likely lead to higher transfers. Future work should explore the influence of this variable on viruses, and specifically non-enveloped viruses, experimentally.

Enveloped virus Phi6 is transferred between surfaces and fingerpads to a lesser extent than non-enveloped virus MS2. This might suggest that enveloped viruses are transferred less efficiently than non-enveloped viruses, however, the effect size is small (difference in mean transfer rate is ~0.1). Both experimental and modeling studies suggest that enveloped and non-enveloped viruses can be transmitted via fomites, and that this transmission requires transfer via a contact event and subsequent self-inoculation. For example, non-enveloped norovirus was shown experimentally in a case study to be transmitted via contaminated surfaces in a houseboat used by different groups in series (22). *Betaarterivirus suid 1*, an enveloped virus that infects pigs, was shown experimentally to be transmitted via contaminated fomites in a controlled animal exposure study (23). Zhao et al. (14) indicate fomites can be important in the spread of enveloped influenza viruses. Boone and Gerba (2) summarize evidence on the role of fomite-mediated transmission of both enveloped and non-enveloped viruses from experimental studies and conclude its role can be important for both types of viruses. It will be important to repeat our study with a broader range of enveloped viruses to confirm the reduced transferability of enveloped versus non-enveloped viruses.

Enveloped virus transfer is higher from smooth plastic and metal surfaces than rough wooden surfaces. Although stainless steel, plastic, and wood are all considered non-porous surfaces, the surface of painted wood is inherently more irregular due to brush strokes. This suggests that the microvariations in the surface of the wood may create a less efficient transfer, and therefore a lower transfer rate of the virus. Such heterogeneities on the surface may prevent efficient contact between fingerpads and the surfaces. Previous studies have modelled that as donor roughness increases, the transfer rate decreases, based on touch probability and adhesive probability (14). However, as recipient roughness increases, the transfer rate correlation is nonmonotonic (14).

Non-enveloped viruses are more readily transferred from surfaces to fingerpads than from fingerpads to surfaces; the mean difference between surface to fingerpad and finger pad to surface transfer rate was found to be 0.23 for plastic, 0.21 for stainless steel, and 0.10 for wood. In previous studies that report that direction of transfer is important in controlling virus transfer, conclusions regarding the direction in which virus was more readily transferred differed based on virus type (5, 10, 12). This agrees with what was found in this study, where only MS2 showed a greater transfer from surfaces to fingerpads than from fingerpads to surfaces. A greater transfer from surfaces to fingerpads than from fingerpads to surfaces suggests individuals are able to pick up viral particles from a surface and may not be able to spread them to additional surfaces as easily. As a result, viruses may remain on the skin rather than be transferred off. Presence of viruses on the hands and subsequent interaction with the nose, eyes, or mouth, may lead to self-inoculation and subsequent infection. A previous study found that the transfer rate for a non-enveloped virus (PRD-1) from fingertip to lip is roughly 34% (21). Additional work investigating skin-to-skin transfer rate, in combination with previous results of surface-to-skin transfer rate, can help develop a complete model of the disease transmission pathway.

We did not find that ‘time since last handwash’ affected transfer of virus between surfaces and fingerpads. In general, handwashing can change the physio-chemical properties of the skin including changing the pH, removing dirt or oil, or leaving behind trace soap chemicals (24). A previous study found that recently washed hands led to decreased transfer of non-enveloped viruses to and from fingerpads and glass and speculated this was a result of changes in moisture level, pH of skin, and other residual effects from the soap (10). Future work that investigates the effects of handwashing under different realistic scenarios, for example with hands that are unwashed for longer periods of time after work outdoors or shopping, may provide additional insights into whether hand washing reduces or facilitates virus transfer between fingerpads and surfaces. It is well understood that hand washing can remove viral pathogens from hands which serves to interrupt transmission pathways involving hand contacts (25).

There are several limitations to this study which have not already been mentioned. First, this study controlled contact pressure even though it is understood that this may affect transfer (12, 14). Additional work to include contact pressure as a variable may be useful. Second, this study worked with clean surfaces and relatively clean fingerpads. In reality, surfaces and fingerpads may be coated with dirt or oil and this could affect transfer rates by changing physio-chemical interactions between viruses and surfaces (14). Further work should consider the use of realistically soiled surfaces and hands, which may provide protection to pathogens when the contact event occurs (26). Third, this study was restricted to two viruses and three surfaces. It would be interesting to expand on these in future studies to investigate whether the trends observed here for enveloped viruses can be confirmed with other surrogate, non-pathogenic enveloped viruses. Finally, our surface sampling technique may not recover all viruses from the surfaces swabbed. An inherent assumption in this work is that the recovery efficiency of virus from fingerpads and tested surfaces was not distinct, so that the transfer rate could be calculated without accounting for recovery efficiency (as recovery efficiency would cancel in the numerator and denominator of Equation 1). Recent work attempts to more accurately represent bacterial concentrations on surfaces using a sequential sampling method (14, 27). Future work should investigate the usefulness of this method for viruses and how its use might affect the calculation of transfer efficiencies.

## Materials and methods

### Volunteers

Volunteers for this study were enrolled with approval from the Stanford University Research Compliance Office for Human Subjects Research according to IRB-55010. 15 volunteers participated per surface, similar to the number of volunteers used in previous studies on virus transfer (10, 18). All volunteers were allowed to participate in the study if they self-reported as healthy, had no visible sores on their hands or fingerpads, and had appropriate building access according to Stanford’s COVID-19 Research Recovery Plan. The experiments were conducted in a room isolated from others, a 6-foot distance was maintained whenever possible, and facial masks were worn at all times according to Stanford’s COVID-19 Research Recovery Plan. Once volunteers were informed of the risks of the experiment and consented, the age, gender, hand length, and hand breadth of the volunteers were recorded. Hand length and breadth was recorded according to the National Aeronautics and Space Administration (28). The volunteer group consisted of 20 volunteers, 8 of whom self-identified as cis-male and 12 as cis-female. Of the 20 volunteers, 9 performed the experiment with all three surfaces, 8 with two surfaces, and 2 with just one surface.

### Virus preparation

Phi6 and MS2 were applied to the surfaces and fingerpads together in the same aliquot to ensure viruses were suspended in equivalent aqueous matrix. An equivalent aqueous matrix is vital to ensure homogenous transfer conditions between the two viruses so that the effect of virus type can be deduced from the experiments. Each virus was diluted to the preferred titer with TSB and then the mixed in equal proportions. TSB was used as the matrix for the experiments to mimic an organic-rich media which better resembles bodily excretions like mucus, saliva, vomitus, and feces than a buffer or water solution.

Phi6 (NBRC 105899) and its host *Pseudomonas syringae* (*P. syringae*, ATCC#21781) were obtained from the University of Michigan. To propagate *P. syringae*, 30 mL of nutrient broth (described in the SI) was inoculated with a loop of *P. syringae* stock from −80°C and incubated while shaking at 30°C for 48 hours until experiment use. The propagated host was kept at 30°C and used for additional experiments up to 48 hours after initial use. Phi6 virus stock was created using the method described in the Supplemental Information (SI) following Wolfe et al. (18).

MS2 (DMS No. 13767) and its host *Escherichia coli* (*E. coli*, DMS No. 5695) were purchased from DSMZ German Collection of Microorganisms and Cell Cultures. 20 mL of tryptic soy broth (TSB, pH of 7·3±0.2) was inoculated with 20 μL *E. coli* stock from −80°C and then incubated at 37°C until the growth phase was logarithmic (about 6 hours), then it was used immediately for experiments. MS2 virus stock was created using the method described in the SI.

### Surface preparation

Samples of the three surfaces were obtained from Home Depot (East Palo Alto, CA, USA). Stainless steel and plastic were light switch cover plates, while painted wood was poplar cut to approximately the same size as the light switch cover plates and painted with interior acrylic semigloss paint (Figure SI.1). 2 cm squares were delineated on the surfaces using permanent marker. To sterilize each surface, the surface was washed with antibacterial soap, soaked in a 10% bleach solution, triple rinsed with DI water, and dried with a Kleenex scientific cleaning wipe (Kimberly-Clark, Irving, TX, USA).

### Experimental protocol

#### Overview

The experimental design of this study was modified from Julian et al. (Figure 2) (10). The experiment can be broken down into two parts, Experiment A and Experiment B. The experiments have the same setup but differ in the length of time since last handwash. Experiment A took place an hour after the volunteer washed their hands with soap and water, while Experiment B took place immediately after handwashing. In both experiments, a donor surface, which represents the contaminated surface, was inoculated with the viruses and the virus inoculum was allowed to dry to mimic the desiccation that can occur during natural contamination events. The donor surface could be one of the three non-porous surfaces tested or could be a fingerpad depending on the direction of transfer. Depending on the volunteer’s schedule, with some volunteers an additional second surface was tested immediately after the first. In all instances, the contact event then took place with the recipient surface (the clean surface(s) or fingerpad depending on the direction of transfer). Samples were recovered from both the donor and recipient surfaces. After Experiment A, the volunteer washed their hands using the same technique as in the beginning of the study, and immediately Experiment B took place. After Experiment B the volunteer washed their hands a final time and the experiment concluded.

#### Detailed Experimental Protocol

A 2 cm × 2 cm square of donor surface (steel, plastic, wood, or fingertip) was inoculated with 10 μL of pooled virus stock containing both MS2 and Phi6. Virus stock consisted of TSB with ~10^5^ PFU MS2 /mL and between 10^8^ PFU Phi6/mL and 10^10^ PFU Phi6/mL. The higher Phi6 titer stock was used for fingerpad and painted wood donor surfaces while the lower Phi6 stock was used for stainless steel and plastic surfaces. The different Phi6 titers were required to obtain countable plaques from the recipient surfaces. Temperature and relative humidity of the room during the experiment were recorded using a ThermoPro TP49 Digital Hygrometer.

An hour prior to Experiment A, volunteers were asked by the technician to wash their hands with antibacterial liquid hand soap (Colgate-Palmolive, New York, NY, USA) for 15 seconds, rinse them in tap water, and dry them with a Kleenex scientific cleaning wipe (Kimberly-Clark, Irving, TX, USA). They were asked to refrain from using the restroom, eating food, and wearing latex gloves until the start of the experiment. For each volunteer, one surface to be tested was chosen through a random number generator from 1-3 (1=Stainless steel, 2=Plastic, and 3=Painted wood). An optional second surface to be tested the same day was also randomly chosen from the remaining 2 surfaces. Next, the finger corresponding to each direction of transfer and the finger used as a control were chosen through a random number generator from 1-5 (1=Thumb, 2=Index, 3=Middle, 4=Ring, 5=Pinky). With each volunteer, one finger served as a recipient for the chosen surface (surface-to-fingerpad transfer), one finger served as a donor for the chosen surface (fingerpad-to-surface transfer), one finger served as a recipient for the second optional surface (surface-to-fingerpad transfer), one finger served as a donor for the second optional surface (fingerpad-to-surface transfer), and one finger served as a control (Figure 2). Collection of control samples, where the virus was not applied to the finger, ensured that there were no viruses present on the hand or surface, and no cross-contamination present. The right and left hands served as duplicates of one another, and as a result the designations were identical for each hand (Figure 2). The viruses were distributed on both the appropriate surface and fingerpads in a grid of small dots (about 0.75 μL per dot) for even distribution and were allowed to visibly dry. This grid was adjusted for each finger, as they had unique sizes, but was approximately a 4×4 grid for surfaces. For surfaces, the drying time typically took about 30 minutes, while for fingerpads it took about 5 minutes.

After the inoculum on the donor surface was visibly dry, the contact event took place. The volunteer contacted the surface for 10 s at a pressure of 25 kPa. The appropriate pressure was administered using a triple-balance beam set to 500 g. This pressure is comparable to a child gripping an object, the pressure of adult fingerpads exerted locally on a hand tool, and studies examining transfer of soil from surfaces to skin (29–31). Upon completion of the contact event, a cotton swab wetted with TSB was used to remove the virus from both the donor and recipient surfaces. The swab was swiped firmly across the surface for 10 s using a sweeping motion. The swab was then placed in 1000 μL of TSB and vortexed for 10 s.

After Experiment A was complete, the volunteer was asked to use alcohol-based hand sanitizer (ABHS) and then wash their hands using the same method they used at the start of the experiments. Immediately after washing, Experiment B was initiated using the same surface(s), and the same fingerpad donor/recipient designations as Experiment A. Experiment B was carried out in the exact same manner as Experiment A. At the end of Experiment B, volunteers were asked to use ABHS again and to wash their hands a final time.

After the volunteer left the experiment, the samples were vortexed, diluted 1:10 and 1:100 using TSB, and then stored at 4°C for a maximum of 8 hours until the plaque assays were performed.

### Quantification

To enumerate Phi6 and MS2 in the samples, traditional double agar plaque assays were used. The Phi6 plaque assay followed Wolfe et al. (see SI) (18). Briefly, soft agar (0.3% agar) was inoculated with 100 μL of sample and 100 μL of *P. syringae* host, then the mixture was poured onto hard agar plates (2.3% agar). The MS2 plaque assay is based on EPA method 1602 (32). Briefly, soft agar (0.7% agar) was inoculated with 300 μL of sample and 200 μL of *E. coli* host, then pouring the mixture onto hard agar plates (1.5% agar).

Three dilutions of each sample were assayed, including undiluted, 1:10 dilution, and 1:100 dilution samples. In addition, a negative control for each hand and surface was included for each volunteer. The negative control consisted of performing the contact event with a surface and fingerpad that were not inoculated with the virus, swabbing the recipient surface, and processing the swab sample using the plaque assay described. The viral stock concentration was enumerated in each experiment, confirming the plaque assay was working correctly even if no plaque were observed in the surface transfer results. The Phi6 and MS2 hard agar plates were incubated at 30°C and 37°C, respectively, for 18 hours before plaques were counted as PFUs. The number of PFUs were counted if the number was between 1 and 500. If there were more than 500 PFU, TNTC (too numerous to count) was recorded. If there were no PFU, then a 0 was recorded.

### Data Analysis

The transfer rate was calculated using the Equation 1. In this equation, the transfer rate (*r*) is defined as the mean PFU times the appropriate dilution factor measured on the recipient surface (*R_R_*) divided by the sum of the mean PFU times the appropriate dilution factors recovered from both the recipient surface and donor surface (*R_D_*). Dilution factor is defined as 1 for undiluted sample, 0.1 for 1:10 diluted, and 0.01 for 1:100 diluted samples. The recovered PFU times the dilution factor was used in the denominator rather than the applied concentration, as desiccation results in a loss of viral titer (10) and we sought to quantify transfer specifically without considering effects of desiccation:

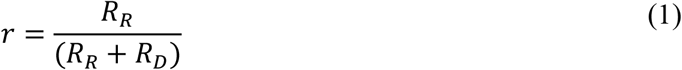

A sample is defined as an individually collected swab of the virus. Each contact event results in two samples, one from the finger swab and one from the surface swab. There are two levels of replication when quantifying the samples for each of the 15 volunteers. The first are the biological replicates created by the duplicate hand profiles of each volunteer. The second are the technical replicates created from the multiple dilutions of each sample. For the purpose of the data analysis, no separation of the biological replicates was attempted. All available technical replicates were multiplied by their appropriate dilution factors and averaged to obtain one recovery from the recipient surface and one recovery value from the donor surface. These are the values then used in Equation 1. Inclusion of the technical replicates can be approached in many ways other than the one chosen (such as only choosing the dilutions that yielded the lowest transfer rate or only using dilutions between a certain range of PFU). Different approaches were tried in the data analysis and no differences in results was noted (details not shown).

Data cleaning and the calculation of the transfer rate was performed in MATLAB (MATLAB R2020a; The MathWorks; Natick, United States). If the PFU count was recorded as TNTC or 0 for either the donor or recipient surface, the data for the transfer event was removed. Descriptive statistics (mean, median, and standard deviations) and statistically modeling functions were calculated in R (R: A Language for Statistical Computing, version 1.2.5042; R Foundation for Statistical Computing, Vienna, Austria). Beta distributions were fit to the data using a univariate maximum likelihood estimation. The goodness of fit was determined through Kolmogov-Smirnoff tests. An n-way ANOVA was used to test the hypotheses that virus type, surface type, time since last handwash, and direction of transfer were significant experimental factors of the virus transfer rate. The n-way ANOVA was followed by a Tukey Honestly Significant Difference post-hoc test. ANOVA assumption testing (including blocking and homoscedasticity) is contained in the SI. A significance level of α=0.05 was used in this assessment.

## Acknowledgments

This work was supported by NSF RAPID (CBET-2023057) to A.B.B. and an NSF-GRF to C. A. We acknowledge the volunteers without whom this study would not have been possible. We also thank Krista Wigginton, Stephanie Loeb, and Marlene Wolfe for their help with virus propagation protocols, and members of the Boehm lab who reviewed and edited this paper. This study was performed on the ancestral and unceded lands of the Muwekma Ohlone people. We pay our respects to them and their Elders, past and present, and are grateful for the opportunity to live and work there.

